# Salp blooms increase carbon export 5-fold in the Southern Ocean

**DOI:** 10.1101/2022.02.07.479467

**Authors:** Moira Décima, Michael R. Stukel, Scott D. Nodder, Andrés Gutiérrez-Rodríguez, Karen E. Selph, Adriana Lopes dos Santos, Karl Safi, Thomas B. Kelly, Fenella Deans, Sergio E. Morales, Federico Baltar, Mikel Latasa, Maxim Y. Gorbunov, Matt Pinkerton

**Affiliations:** National Institute of Water and Atmospheric Research (NIWA), Hataitai, Wellington 6021, New Zealand; Department of Earth, Ocean, and Atmospheric Science, Florida State University, Tallahassee, Florida, 32304, USA; Center for Ocean-Atmospheric Prediction Studies, Florida State University Tallahassee, Florida, 32310 USA; Department of Oceanography, University of Hawai’i at Mānoa, Honolulu, Hawaii 96822, USA; Asian School of the Environment, Nanyang Technological University, 50 Nanyang Avenue, Singapore 639798; National Institute of Water and Atmospheric Research, P.O. Box 11-115, Hamilton, New Zealand; Department of Microbiology and Immunology, University of Otago, Dunedin, New Zealand; Department of Functional & Evolutionary Ecology, University of Vienna, 1090, Austria; Instituto Español de Oceanografía, Centro Oceanográfico de Gijón, Avenida Príncipe de Asturias, 70 bis, 33212 Gijón, Spain; Environmental Biophysics and Molecular Ecology Program, Department of Marine and Coastal Sciences, Rutgers, The State University of New Jersey, New Brunswick, NJ 08901 USA

## Abstract

The Southern Ocean (SO) contributes substantially to the global biological carbon pump (BCP). Salps in the SO, in particular *Salpa thompsoni*, are keystone grazers that produce large, fast-sinking fecal pellets with high export potential. In a first study of this kind, we conducted Lagrangian experiments to quantify the salp bloom impacts on export pathways by contrasting locations differing in salp bloom presence/absence. We show that blooms increased particle export by ~5-fold, and exported up to 46% of net primary production out of the euphotic zone. BCP efficiency increased from 5% in non-salp areas to up to 28% in salp areas, which is among the highest recorded in the global ocean. Using SO salp abundances from KRILLBASE, we estimate they can consume ~ 13% of regional production, mediating 13-40% of the SO BCP. Consideration in models forecasting the SO BCP is recommended considering long-term increases in SO salp abundances.

---

The Southern Ocean (SO), which covers ~1/3 of the world’s ocean, plays a fundamental role in global climate. The biological carbon pump (BCP), defined as ecosystem processes that transport organic carbon to depth, has profound implications at planetary scales, from the architecture and efficiency of food webs to the sequestration of carbon to deep waters. The SO BCP accounts for a significant (2-3 Pg C y^−1^) fraction of the global BCP (5-13 Pg C y^−1^)^1^. Distinct biomes are found in the SO with variable efficiencies in BCP strength^2^. Predicting the response of the SO BCP to climate change requires elucidation of ecological mechanisms that drive BCP variability^3^ in this important ocean area.

Salps are gelatinous zooplankton grazers that are widespread in the SO^4^. Climate change-induced ocean warming may be allowing salps to expand into colder SO waters in the vicinity of the Antarctic continent, where they can potentially displace krill (a keystone species)^5^. The southward range expansion of salps may have important consequences for the BCP, and biogeochemical models will benefit from a quantitative understanding of their effects on the marine carbon cycle. Substantial salp contributions to the BCP have long been hypothesized.

High ingestion rates^6^, extensive bloom formation^7^, ability to feed on a wide range of prey sizes (1-1000μm)^8,9^, and fast fecal pellet (FP) sinking speeds^10^ of large pellets^11^, hint at the potential of salps to enhance ocean carbon sequestration^12^. However empirical evidence quantifying the effect of salp blooms on carbon export budgets is very limited, mainly because it requires simultaneous measurements of other plankton-mediated processes controlling export. Models suggest substantial contribution of salps to the BCP^13,14^, but lack data necessary for validating their many assumptions. While the episodic nature of salp blooms has precluded controlled studies in the past, we successfully predicted the timing and location of salp blooms in the SW Pacific sector of the Southern Ocean, and present results from the first plankton food-web process study quantifying salp-bloom impacts on BCP efficiency.

Our oceanographic voyage, SalpPOOP (Salp Particle expOrt and Ocean Production), was conducted during austral spring (October/November 2018) in SO waters near the Chatham Rise (Fig. 1a), where nitrogen-limited Subtropical (ST) and iron-limited Subantarctic (SA) surface water masses meet to form the circumglobal Subtropical Front (STF)^15^. Our experimental design featured Lagrangian experiments following water parcels in ST and SA waters for a period of 3-7.5 days (Table 1). We conducted repeat measures of hydrographic and nutrient conditions, plankton rates and stocks, and assessed particle export flux using free-drifting Particle-Interceptor Traps (PITs) at different depths (typically 70, 100, 300 and 500m) and ^238^U-^234^Th disequilibrium methods^16^. To test the hypothesis that salp FPs enhance the BCP, we investigated flux patterns in five experimental locations, within two distinct water types of the SO, and in water parcels with and without salp blooms. This regional study was put into a broader context by using our relationships between abundance, grazing, and carbon export to obtain an estimate of the possible impact of salps on primary production and export flux within the SO^17^.

**Table 1.** SalpPOOP Experiment cycle characteristics. Cycle duration, nutrient (nitrate and silicate) concentrations, areal net primary production (NPP) and chl *a* integrated for the euphotic zone (0.1% surface PAR), non-salp zooplankton biomass (to 200m), non-salp zooplankton grazing, salp abundance, and salp composition. Salp SA-Sc had 2 deployments of Particle Interceptor Traps (PIT), so the cycle is divided into A and B time-periods. Experiments are listed in the order they were sampled. Note that the most abundant salp was *S. thompsoni*, and the abundances of the ‘other salp species’ are aggregated, since they were low, to indicate their contribution to both ‘salp’ and ‘non-salp’ locations (mean ± SE).

| Cycles in order sampled |  | Days | Nitrate (mmol m<sup>-2</sup>) | Silicate (mmol m<sup>-2</sup>) | NPP (mg C m<sup>-2</sup> d<sup>-1</sup>) | Areal chl *a* (mg m<sup>-2</sup>) | Zooplankton (mg DW m<sup>-2</sup>) | Zooplankton grazing (mg chl *a* m<sup>-2</sup> d<sup>-1</sup>) | *S. thompsoni* (salps m<sup>-2</sup>) | Other salps (salps m<sup>-2</sup>) | Other salp species |
| --- | --- | --- | --- | --- | --- | --- | --- | --- | --- | --- | --- |
| Salp SA-Sc | A | 5 | 379 ± 81 | 58 ± 7 | 638 ± 236 | 39 ± 4.8 | 2.1 ± 0.6 | 4.9 ± 0.8 | 831 ± 313 | 3 ± 3 | *Soestia zonaria* |
| | B | 2.5 | 326 ± 40 | 40 ± 14 | 387 ± 10 | 49 ± 5.1 | 2.3 ± 0.8 | 3.5 ± 0.6 | 1649 ± 656 | 1 ± 0.5 | *Soestia zonaria* |
| Salp SA | | 5.5 | 527 ± 151 | 43 ± 15 | 321 ± 62 | 28.1 ± 1.9 | 1.6 ± 1 | 1.2 ± 0.2 | 212 ± 54 | 4.3 ± 2.2 | *Thetys vagina*, *Pegea confederata* |
| Non-Salp ST | | 3 | 57 ± 19 | 12 ± 5 | 985 ± 107 | 64.9 ± 9.6 | 3.6 ± 1.7 | 1.2 ± 0.1 | 0 | 6 ± 3 | *Soestia zonaria*, *Salpa fusiformes*, *Ihlea magalanica*, *Thalia democratica* |
| Salp ST | | 4 | 146 ± 29 | 4 ± 1 | 452 ± 102 | 46.5 ± 12.3 | 1.8 ± 0.7 | 1.1 ± 0.1 | 70 ± 15 | 2 ± 1 | *Soestia zonaria*, *Salpa fusiformes*, *Ihlea magalanica*, *Thalia democratica* |
| Non-Salp SA | | 3 | 820 ± 167 | 58 ± 20 | 251 ± 30 | 26.5 ± 10 | 0.9 ± 0.6 | 0.5 ± 0.1 | 0 | 1 ± 0.3 | *Soestia zonaria* |

**Fig. 1.**
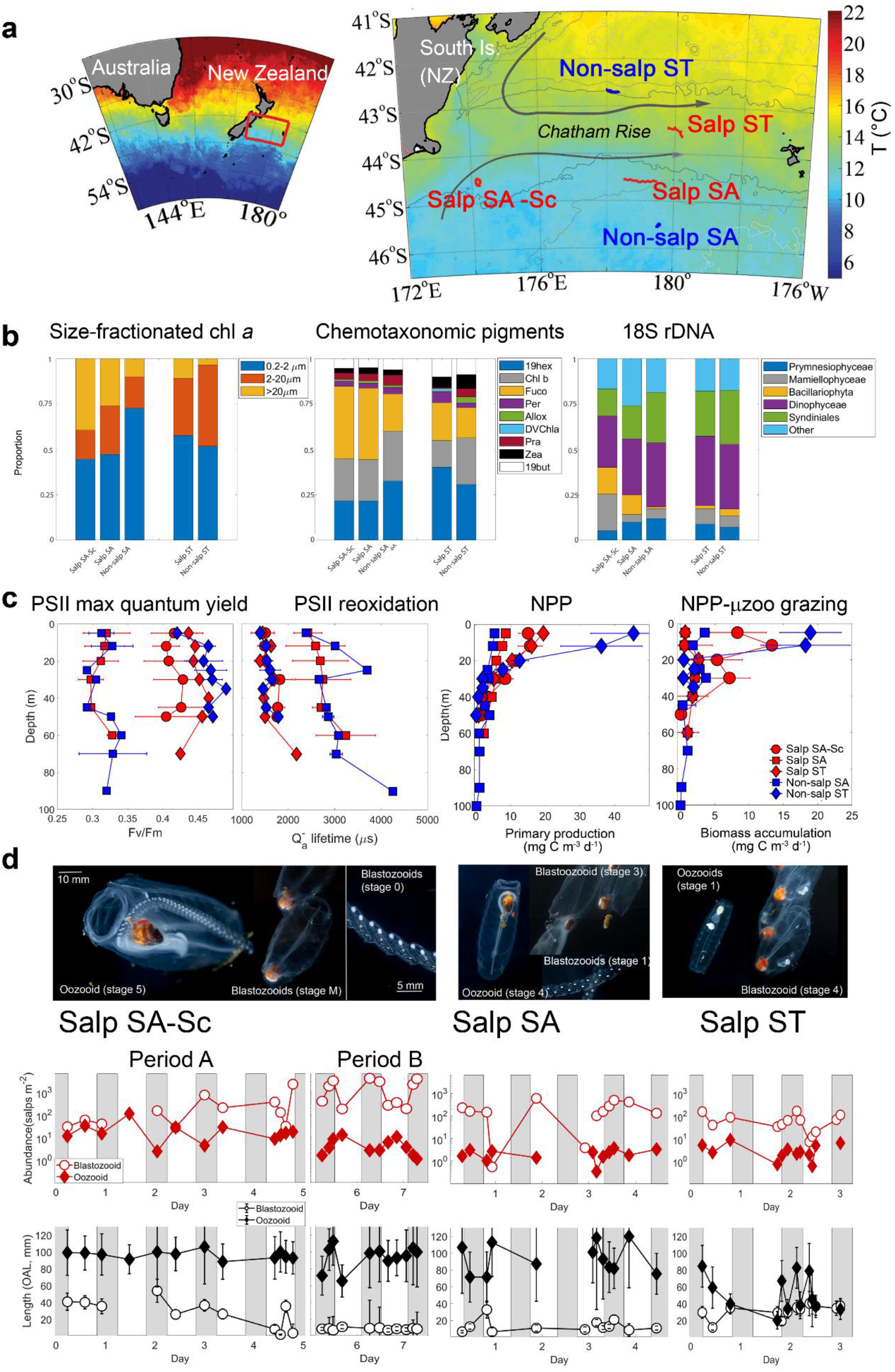
Study area, phytoplankton community, and *S. thompsoni* bloom composition. (**a**) Study area, currents, and Lagrangian sampling (tracks shown as red or blue lines/dots). Water mass types: Subantarctic Southland Current (SA-Sc), Subantarctic (SA), and Subtropical (ST). (**b**) Autotrophic integrated community composition: size-fractionated chlorophyll *a*, pigment-based composition, and DNA-based composition. (**c**) Phytoplankton physiology indicated by PSII maximum quantum yield (Fv/Fm) and reoxidation kinetics (Qa—lifetime), net primary productivity (NPP) and phytoplankton biomass accumulation (NPP – μzoo grazing). Red markers indicate Salp bloom locations, and blue markers indicate Non-salp locations. Circles and squares = SA waters, diamonds = ST waters (**d**) Abundances (salps m^−2^) and mean (± weighed standard deviation) OAL for blastozooids (aggregates) and oozooids (solitary) of *S. thompsoni*. Note Salp SA-Sc was divided into Period A (first 5 days) and Period B (subsequent 2.5 days). Grey bars indicate night times, white bars indicate day times. Images are from each sampling location, indicating the predominant stages sampled: early bloom for Salp SA-Sc, intermediate for Salp SA, and late bloom stage for Salp ST^21^.

## RESULTS

### ST and SA water masses: conditions and salp bloom evolution

The problem of finding salp blooms to conduct this study was addressed by capitalizing on knowledge of ‘salp-eating’ New Zealand deep-sea fishes: oreos (Oreosomatidae) and warehous (*Seriorella* spp.)^18,19^. Their well-defined demersal habitat areas over the Chatham Rise^18,19^ were successfully used to target our salp-bloom search. Five water parcels (hereafter referred to as ‘cycles’) were investigated. This included three SA cycles (one influenced by the Southland Current, which is a primarily (90%) SA current ^20^ [designated as ‘SA-Sc’], and two in SA proper) and two ST cycles (Fig. 1a, Supplementary Fig. 1&2). One cycle in each water mass did not have a salp bloom, for a controlled comparison with conditions where salps predominated in the same water mass. Temperature-salinity (T-S) data supported the classification of water parcels as SA or ST, although some mixing was evident in the T-S characteristics of Salp SA-Sc and Salp ST sites (Supplementary Fig. 2). Phytoplankton size, pigment and DNA-based water community composition also supported the classification of the water parcels as SA and ST (Fig. 1b). Phytoplankton cells larger than 20μm were more abundant in SA waters, especially close to the Chatham Rise, while a greater nanophytoplankton (2-20 μm) contribution was observed in ST waters (Fig 1b, left panel, Supplementary Fig. 3). In general, diatoms, represented by Fucoxanthin (Fig. 1b, middle panel, Supplementary Table 1) and Bacillariophyta 18S rDNA abundance (Fig. 1b, right panel), were more abundant in SA while ST was dominated by Dinophyceae (Fig 1b, right panel). Phytoplankton physiology measured using variable fluorescence and the kinetics of Photosystem II (PSII) reaction centers suggested iron-stress in phytoplankton inhabiting SA waters, but Salp SA-Sc had physiological conditions similar to ST, indicating that frontal mixing alleviated nutrient stress (Fig. 1c). Net primary production (NPP) was highest in ST waters, intermediate for Salp Sa-Sc and lowest for SA waters (Table 1).

Phytoplankton biomass accumulation – i.e. phytoplankton that escape microzooplankton grazing and can be potentially exported through direct sinking – was also highest in Non-salp ST, followed by Salp SA-Sc (Fig. 1c). Salp blooms were dominated by *Salpa thompsoni*, with distinctly different age and stage structures in different locations^21^ (Fig. 1d, Table 1). The Salp SA-Sc cycle was characterized by an early-stage bloom. This experimental cycle lasted 7.5 days, although PIT arrays were recovered on day 5 and re-deployed for an additional 2.5 days, so we divide the experiment into 2 periods (A, B). This cycle exhibited the highest abundance of salps, including initial high abundances (10-100 individuals m^−2^) of the large oozooid (asexual, solitary) stage (Period A). Small blastozooid (sexual, aggregate) abundance increased over the later part of the experimental cycle (up to 1512 ± 456 ind. m^−2^ (mean ± SE), Period B). Salp SA exhibited an intermediate bloom stage, with larger blastozooids and the lowest oozooid abundance of all locations (2.0 ± 0.3 individuals m^−2^). Finally, Salp ST had larger blastozooids and younger, smaller oozooids, indicating the completion of their reproductive cycle (Fig. 1d)^21^. Although salps were detected in the non-salp locations, these were often other species than *Salpa thompsoni* (Table 1) and in very low abundances.

### Salp grazing and fecal pellet contribution to the BCP

Salp bloom conditions were characterized by much higher particulate organic carbon (POC) export than control (non-salp) locations (Fig. 2a), particularly in ST waters. During salp cycles, POC fluxes below the euphotic zone ranged from 80 to 210 mg C m^−2^ d^−1^ in surface waters (Salp-SA = 80-130 mg C m^−2^ d^−1^, Salp-ST = 210 mg C m^−2^ d^−1^). Export into the mesopelagic zone at 300-m depth ranged from ~55 mg C m^−2^ d^−1^ (Salp-SA-SC= 43 ± 7 mg C m^−2^ d^−1^; Salp-SA= 59 ± 18 mg C m^−2^ d^−1^) to ~120 mg C m^−2^ d^−1^ (Salp-ST= 119 ± 49 mg C m^−2^ d^−1^). Particle export in non-salp areas was significantly lower (Non-salp ST= 34 ± 2 mg C m^−2^ d^−1^; Non-salp SA = 25 ± 12 mg C m^−2^ d^−1^ below the euphotic zone; mesopelagic flux (300m) values decreased to Non-salp ST = 19 ± 6 mg C m^−2^ d^−1^; Non-salp SA = 7 ± 6 mg C m^−2^ d^−1^) (Fig. 2a). Particle flux measurements from independent ^238^U-^234^Th disequilibrium approaches generally agreed with the results from the PIT sediment trap arrays, both in terms of overall magnitude of sinking POC and in the relative flux patterns between cycles (Supplementary Fig. 4). However, lack of steady state was clear for Salp-SA, where export estimates were an order of magnitude higher when non-steady state equations were used (Supplementary Figure 4). In addition, Non-salp SA had disproportionally higher ^238^U-^234^Th – based export, which was likely due to a prior high flux event that depleted ^234^Th from the water column (not shown). Capturing the effects of the episodic highly variable nature of salp blooms using this method is complicated due to the difference in time-scales of the event and ^238^U-^234^Th integration. Given that the ^238^U-^234^Th – disequilibrium method is logistically the easiest method of estimating export (requiring only CTD water in contrast to following sediment traps) and thus most commonly used, it is important to understand the strengths and limitations in estimating carbon flux to depth. Our results support its use for general regional estimates of export, but indicate limitations in detecting short-term episodic events, in particular in highly advective areas of the ocean.

**Fig. 2.**
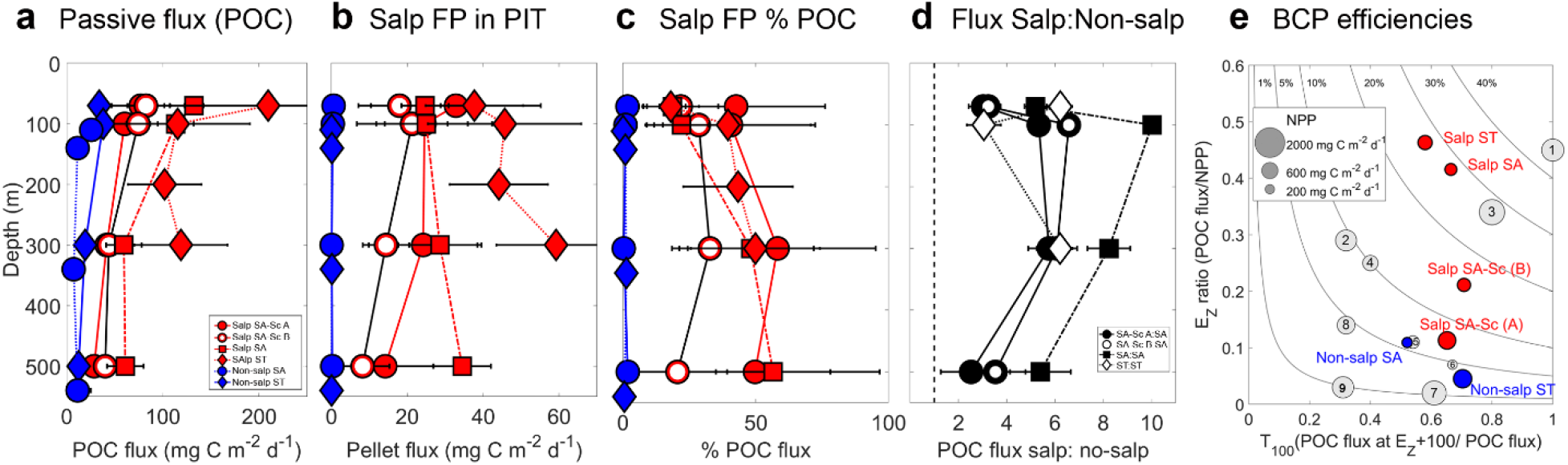
Patterns in carbon export flux. (**a**) Export fluxes of particulate organic carbon (POC), (**b**) Carbon flux due to recognizable salp fecal pellets (FP), (**c**) Relative contribution of salp FP to POC flux, (**d**) Ratio of POC flux between Salp and Non-salp locations, with three comparisons for SA waters, and one for ST waters. (**e**) EZ ratio (POC flux : NPP) as a function of T100 (POC flux at EZ + 100m : POC flux), or flux attenuation. Numbers indicate: 1 – North Atlantic Bloom Experiment (NABE) (spring, temperate North Atlantic), 2 – Kiwi 7, 3 – Kiwi 8 (Polar Front, Pacific sector, Southern Ocean), 4 – K2-D1 (subarctic NW Pacific), 5 – K2 - D2 (subarctic, NW Pacific), 6 – ALOHA (subtropical, central North Pacific), 7 – EqPac (tropical, central Pacific), 8 – OSP – Aug (summer, NE Pacific), 9 – OSP – May (spring, NE Pacific). Circles are proportional to magnitude of NPP (See legend insert). Data from other locations from^2^. Blue indicates non-salp locations, red indicates salp cycles during the SalpPOOP experiment.

Microscopical analyses of PIT contents indicated that in salp areas the contribution of intact salp FPs was significant (Fig. 2b). In salp areas, 20-40% of export at the base of the euphotic zone was directly attributable to recognizable salp FPs in PIT samples (a conservative estimate of their true contribution to flux), and salp pellets accounted for up to 50% of the mesopelagic (300-500m) flux (Fig. 2c). The ratio of total flux in salp:non-salp areas varied with location and depth, but on average salp blooms enhanced the magnitude of carbon export through the water column by ~5-fold (range 2‒10-fold) (Fig. 2d).

The efficiency of the BCP is the product of the proportion of NPP that sinks out of the euphotic zone (EZ) and the proportion of this flux that sinks an additional 100m into the mesopelagic zone (T100, the flux transmission^2^). The EZ ratio was highly enhanced in areas with salps compared to non-salps (Fig. 2e). The EZ ratio also seemed to increase with the stage of the salp bloom (Fig. 2d, Fig. 1d). In ST waters, the EZ ratio was 0.05 without salps and increased to 0.46 with salps; while in SA waters it increased from 0.11 to 0.42 due to the presence of salps. T_100_ varied between 50-70% of POC at the EZ sinking to 100m below the euphotic zone. BCP efficiencies (% of NPP that was exported deeper than 100m below the euphotic zone) for the oceanic salp locations was ~28%, which is among the highest efficiencies recorded in the global ocean^22^ (Fig. 2e).

### Microplankton communities and other potential drivers of carbon export

The effect of microbial communities in regulating export was investigated through community genetic analyses of three different groupings: PIT protists, water column (WC) protists, and WC prokaryotes. Variability between communities was explained primarily by water mass (SA/ST explained 7%, 19% and 20% variance, respectively, for the three groups), but salp/non-salp delineations also explained 3% of variability in PIT and 8% in WC samples (Fig. 3).

**Fig. 3.**
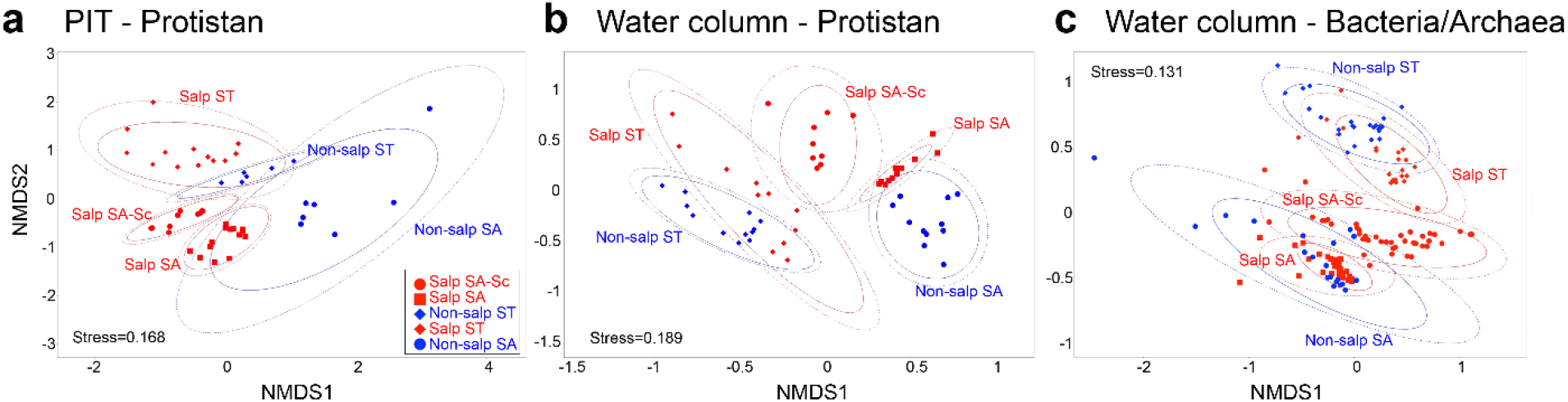
Non-metric multidimensional scaling (NMDS) of microbial communities during SalpPOOP. (**a**) Protistan communities sampled in Particle Interceptor Traps (PITs), (**b**) Protistan communities in the euphotic zone of the water column (WC), sampled with the CTD water bottles, and (**c**) Bacteria/Archaea communities in the euphotic zone (CTD water bottles). Water mass explained most of the variance among communities in the different locations - PIT protists: 7% variance, WC protists: 19%, WC prokaryotes: 20% (PERMANOVA, all *p*-values <<0.001). Salp/non-salp followed for protists (PIT: 4% variance, WC: 9% variance), and was also significant for prokaryotes (7.6% variability). Depth of collection explained similar levels of variability in protistan communities (PIT: 3%, WC: 8%), and was more important for prokaryotic communities (16% variance).

Phytoplankton composition in PITs did not indicate that any one of the main measured algal groups were driving export. Diatoms (bacillariophyta) and coccolithophores (prymnesiophytes) would be expected to drive export flux (due to ballasting), but neither of these groups (or any other) presented in noticeably higher contributions in PIT compared to WC (Fig. 4), and none of these groups were statistically correlated with high export fluxes (Supplementary Table 2).

**Figure 4.**
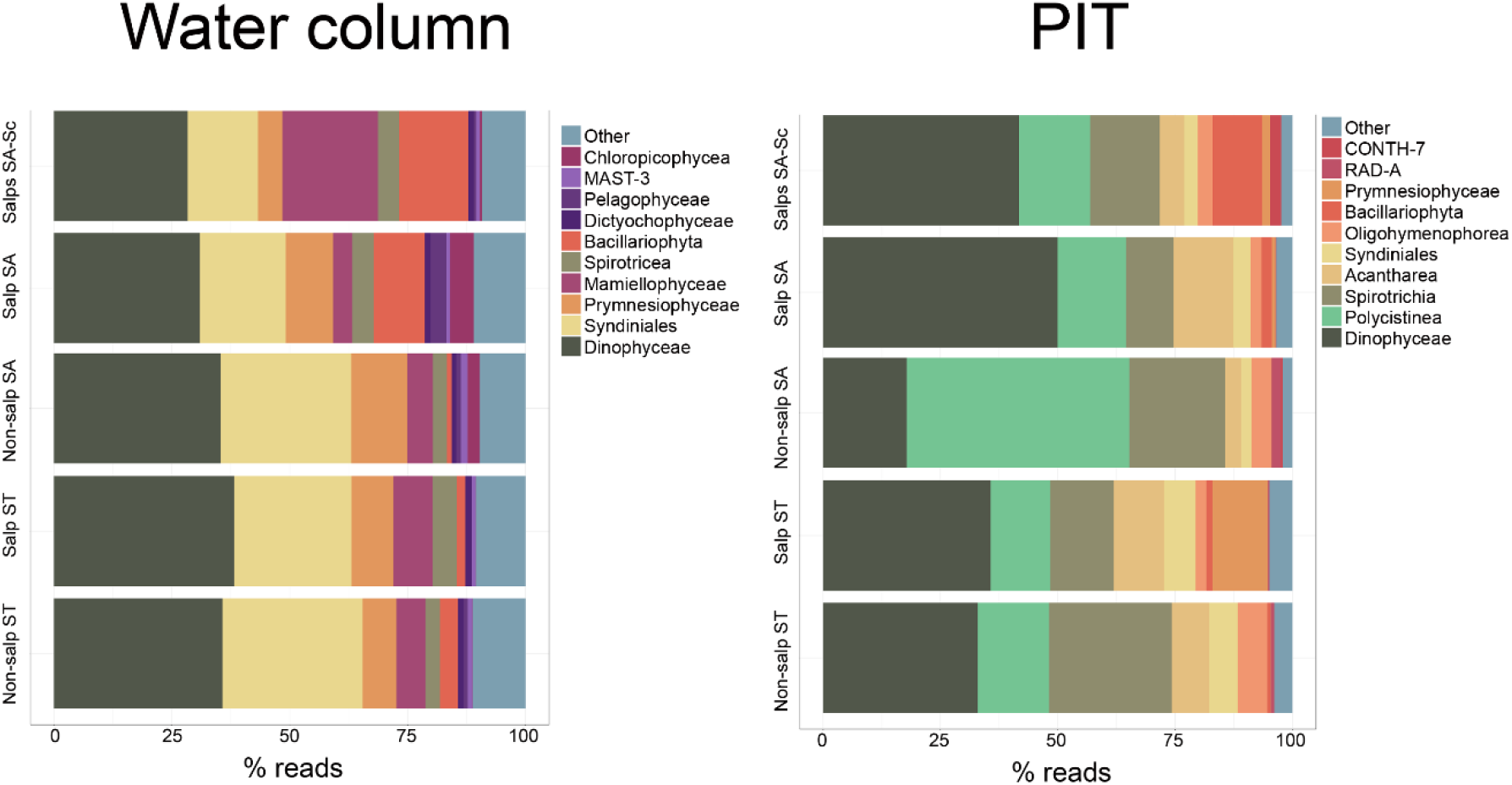
Percent DNA meta-barcoding read contributions of the main 10 plankton groups in Water column and PIT samples. Water column includes six depths spanning the euphotic zone, while PIT sequences are from formalin-fixed sediment trap samples at four depths (see text for details).

### Export estimates derived from salp grazing and potential contribution in the SO

Parallel estimates of salp FP production during SalpPOOP, determined by multiplying Ingestion (grazing rate, Graz) by Egestion Efficiency (EE=30%)^23^, suggested even higher potential FP contributions for SA locations compared to PIT microscopy (Fig. 5). A temporal lag between bloom stage, grazing, and export likely explains patterns across the four locations where we have paired PIT export and salp grazing estimates (3 cycles plus period A and B of Salp SA-Sc). During Salp SA-Sc (A) bloom, Graz×EE exceeded microscopic observations in PITs. This excess increased dramatically during the Salp SA-Sc (B), concurrent with order-of-magnitude increases in salp numbers (Table 1, Fig. 1d). Salp SA had a lower excess of grazing compared to export, and in Salp ST (the area of the most advanced bloom) estimates were in close agreement with PIT microscopy (PIT microscopy = 47 ± 9 mg C m^−2^ d^−1^, cf. Graz×EE = 32 ± 7 mg C m^−2^ d^− 1^). Collectively, the regional averages indicate that during blooms salps mediate 60% (range 20-96%) of POC export at the surface and 50-150% at deeper (>300m) depths.

**Figure 5.**
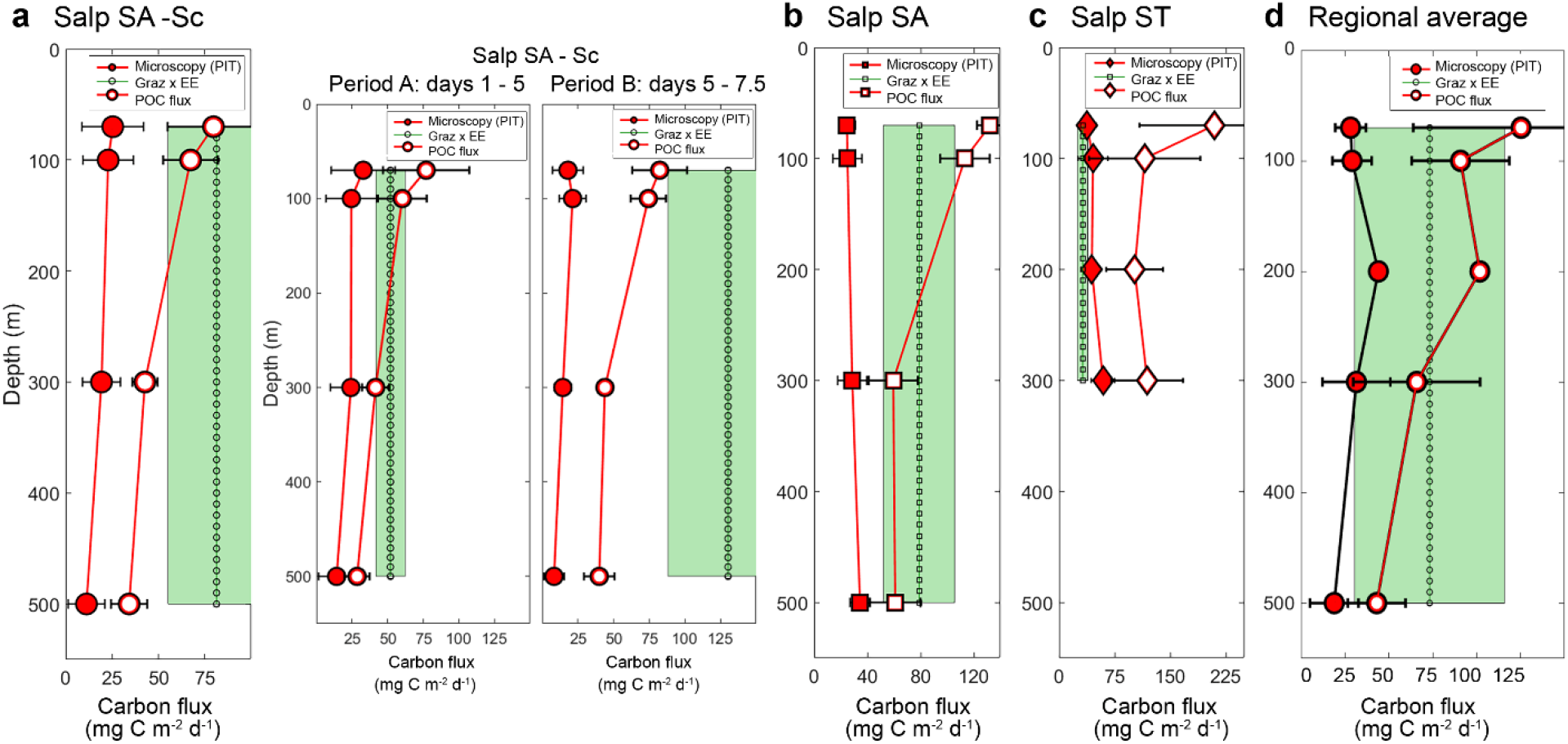
Salp contribution to POC export based on measured grazing (Graz) and egestion efficiency (EE=0.3). Graz×EE, compared to fecal pellets (FPs) (from microscopy of PIT samples) and total POC flux measured in PITs. (**a**) Salp SA-Sc, two subpanels indicating Period A (first 5 days) and Period B (last 2.5 days). (**b**) Salp SA, (**c**) Salp ST, and (**d**) Regional averages of three salp cycles.

Particle export based on sediment trap measurements was thus in line, from a regional perspective, with these estimates based on constraining fecal pellet production from grazing and known egestion efficiencies^24^ – suggesting that on this regional scale all of the FP production estimated based on these egestion efficiencies could be exported, since there is still substantial non-salp mediated flux (40%). Capitalizing on the conclusion that salp grazing is an adequate predictor of salp-mediate export flux, we further developed an empirical relationship with salp abundance. This C-based grazing-abundance relationship (Supplementary Fig. 5) was used to estimate the proportion of SO NPP grazed by salps, and the likely contribution to the BCP in the entire SO since *S. thompsoni* is the dominant salp species in the SO^5^. The C-based grazing estimates with size from our study were comparable to *S. thompsoni* grazing rates measured in polar Antarctic waters^25^. Using satellite-derived NPP estimates^26^ and mean salp abundances from the SO KRILLBASE^17^ (both averaged over the November-March period), we estimated that salps consume on average 380 Tg C y^−1^ in the area south of the Polar Front (Fig. 6), which corresponds to 13% of the regional SO NPP, egesting 3.9 % NPP, assuming an EE of 0.3^24^.

**Fig. 6.**
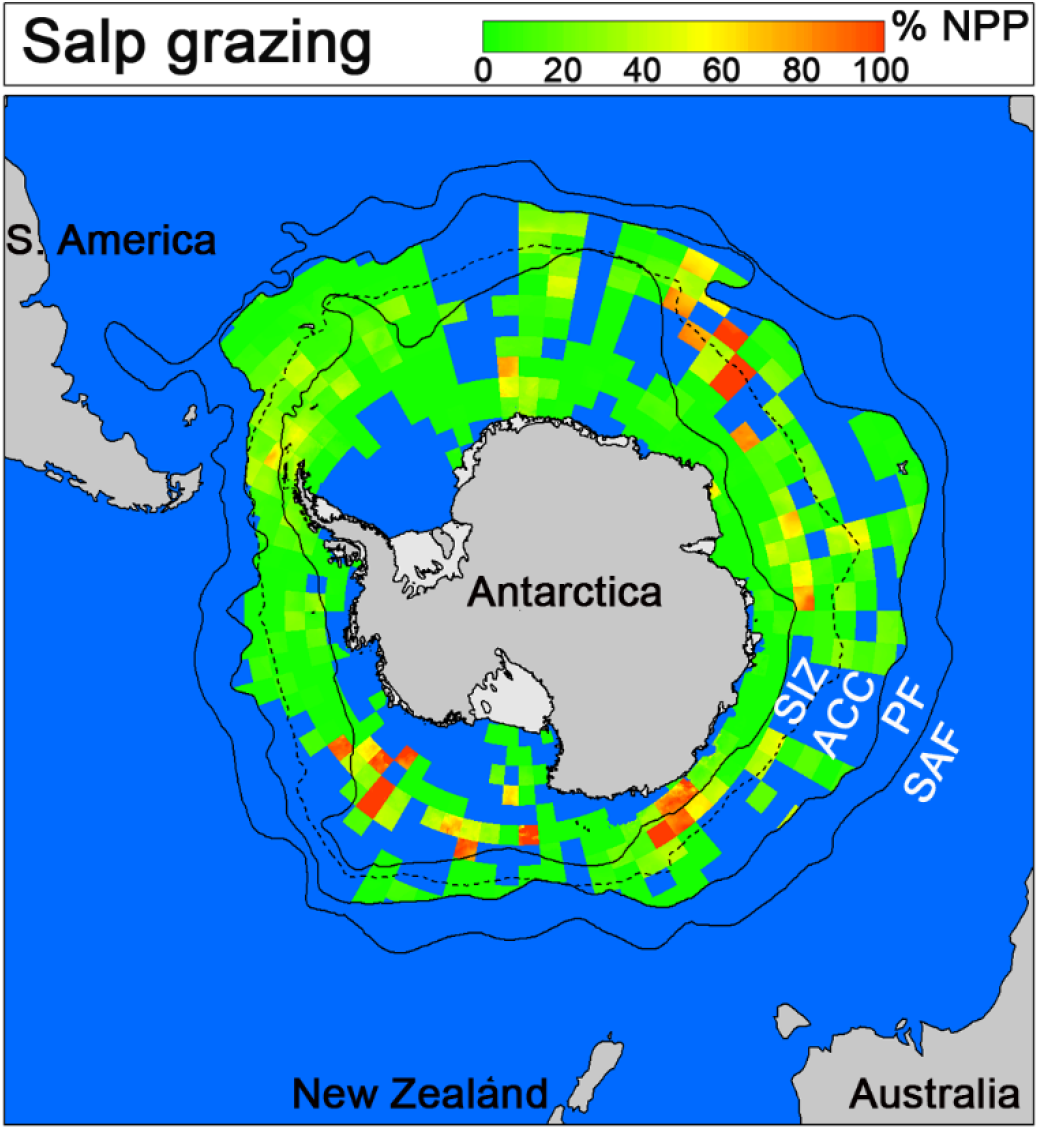
Salp grazing in the SO. Lines indicate the Subantarctic Front (SAF), Polar Front (PF), Southern ACC Front (ACC), and northern sea-ice zone (15% sea-ice concentrations, SIZ). Color is proportional to salp grazing (green – low to red – high), colors indicate % NPP grazed.

## DISCUSSION

Our results demonstrate a pronounced shift in regional POC export fluxes and efficiency of the BCP strongly driven by the presence or absence of salp blooms (Fig. 2a-d). Export in the five areas was largely uncorrelated with nutrient and plankton variables (Supplementary Table 2, Fig. 4) and could not be explained by other potential drivers of the BCP. While presence/absence of salps was the primary determinant of carbon export, total salp biomass was weakly correlated with export, likely because of spatial/temporal decoupling related to the evolution of the bloom. While over any short-period of time (3-5 days) salp grazing could over- or slightly under-estimate salp FPs in PITs (Fig. 5), the regional budget indicated that salp FP production was 20-100% of total export, and within the range of microscopy-determined FP contributions. This supports this approach as a useful proxy in estimating salp contributions to carbon flux. While the regional EZ ratio for the SO is not well constrained, ranging 0.1-0.3 (Fig. 2e, studies 2 and 3) of NPP, salps could be responsible for 13-40% of the SO BCP, assuming (based on this study, Fig. 5) that all of salp egestion (4% NPP) is exported out of the euphotic zone as salp FPs. In addition, because this estimate still only accounted for 60% of total export in salp areas, and flux was always enhanced by at least a factor of 2 (Fig. 2d, 2-8), this seems appropriate as a broad approximation of their effects over the ecosystem in which they are keystone grazers. Contributions to the mesopelagic budget would likely be substantially higher given low flux attenuation of rapidly sinking pellets (Fig. 2d), but this is not well constrained within the SO.

### Decoupling of salp grazing and export

While our study undoubtedly shows that salps enhance export by ~5-fold, the trends in salp grazing and export within salp experimental cycles were somewhat opposite. For instance, in Salp SA-Sc cycle, salp-mediated export estimates from grazing measurements and from microscopical analysis of PIT contents were more similar in the first 5 days as the bloom developed (Graz×EE : FP(PIT) = 2), than the latter 3 days (Graz×EE : FP(PIT) = 8). This increased mismatch was associated with bloom development and a large increase in the number of small blastozooids (producing much smaller pellets) that grazed high amounts of phytoplankton. The Salp SA cycle, with a more advanced bloom, had a value of Graz×EE : FP(PIT) = 2.8, and finally Salp ST had Graz×EE : FP(PIT) = 0.7, despite being the location with the highest export rates. Based on the prevailing patterns of advection, and the dominant salp stages found in each water parcel, Salp ST waters were hypothesized to have the most advanced bloom^21^ we sampled during SalpPOOP. This advanced stage bloom was associated with both higher total export and a higher ratio of PIT-estimated FP flux to grazing than younger stages of the bloom.

In addition to the observed patterns of decoupling in export and grazing, methodological limitations can further complicate attempts to relate these episodic events to export patterns. The lower inter-cycle variability of ^238^U-^234^Th disequilibrium-derived fluxes compared to PIT, and a noticeable discrepancy between the sediment trap and ^234^Th results in the non-salp SA cycle (Supplementary Figure 4), both indicate the limitations related to the longer integration time (approximately one month)^16^ of ^234^Th methods. Export estimated using this method was substantially greater than sediment trap-derived flux in the non-salp SA cycle, likely because of advection of low ^234^Th water (impacted by a prior export event) to the study site. Such events are likely when ST waters mix with SA waters in the vicinity of the STF^15,27^. We found substantial copies of *S. thompsoni* DNA sequences, both in the water and the sediments (not shown), suggesting a prior flux event similar to the types of blooms we were sampling, although by the time we arrived at the site there were no *S. thompsoni* specimens remaining in the water (Table 1). Other studies in the Atlantic sector of the SO, using ^238^U-^234^Th disequilibrium-derived fluxes and short (~1day) PIT deployments may have failed to detect high fluxes specifically associated with salp blooms ^28–30^ because of this spatial and temporal decoupling. The same studies found that, on a regional scale, higher flux transmission to 400m was present in areas south of the PF, where both diatoms and salps were abundant.

Decoupling between production and export have been reported extensively in the literature on regional scales^31–33^, yet the expectation based on fast-sinking of salp FPs was that these would be much more closely linked. On the large scale, they were, as salp locations had 5-fold higher flux. However, the specific details relating flux to salp population grazing supported temporal and spatial lags even when particles are dominated by large, fast sinking FPs.

However, the only other study (to our knowledge) investigating *S. thompsoni* FPs attenuation between 100 and 300m^34^ also found high variability: 20% to 100% of FPs measured at 100m sank to traps at 300m (mean 58% ± 40%^34^), supporting the conclusion that a substantial portion of FPs are either remineralized or sink at slower speeds such that they are not collected during short sediment trap deployments^34^. This study also had a bloom characterized by smaller salps (<20mm) and order of magnitude fewer animals. Smaller pellets and concomitant slower sinking speeds would decrease export efficiency. While the SalpPOOP study provides solid evidence for the role of salps in enhancing export, and our ability to estimate their effect on regional scales (as shown in Fig. 5), our results and these past studies^34^ underscore the complexity of the BCP and the many processes involved in regulating carbon transfer from the surface to the deeper ocean, requiring further study.

While salp locations had enhanced vertical particle flux (Fig. 2d), the magnitude of export within salp cycles was inversely related to salp grazing and positively related to bloom age, with highest export rates in the declining bloom stage of Salp ST (Fig 1d). The lack of a change in T100 flux transmission (which remained between 0.5-0.7, Fig. 2e) between salp/non-salp locations also supports this decoupling - expected values in salp locations would be closer to 1 compared to non-salp areas if pellets sink as quickly *in situ* as they do in laboratory settings.

The SalpPOOP study provides clear evidence for the role of salps in enhancing export. The EZ ratio in both SA/ST waters without salps was similar to traditional low-flux environments such as the North Pacific Subtropical Gyre (e.g. studies 6 and 7 in Fig. 2e). In contrast, salp blooms shifted the ecosystem to conditions comparable to high-flux regions such as the North Atlantic (study 1 in Fig. 2e) following the annual spring bloom and its demise, and the Southern Ocean Polar Frontal Zone during the austral diatom bloom in spring (studies 2 and 3 in Fig. 2e). It should be noted that this study only addresses salp-mediated export in the form of FPs, however export due to sinking carcasses and active diel vertical migration would suggest an overall enhanced effect on the marine C-cycle when all pathways are considered. Finally, investigations of changing SO biogeochemistry often focus on impacts mediated by changing NPP. For instance, a SO export efficiency meta-analysis suggests that a doubling of NPP would increase carbon export by ~50% or less^35^. Our results highlight the importance of factors other than NPP as key players in SO biogeochemistry, revealing that a switch from crustacean- to salp-dominated communities could lead to a 200% - 1000% enhancement of POC export. While a region-wide shift to complete salp-dominance in the SO should not be expected, future studies should now determine if the current long-term increasing trend in salp abundance (+66%)^5^ is widespread through the SO, whether this is likely to persist under future warming conditions^36^, and explicitly include salps in regional ecosystem and biogeochemical models.

## Materials and Methods

### Oceanographic sampling

Sampling for SalpPOOP was carried out onboard the *R/V* Tangaroa from October 23^rd^ to November 21^st^, 2018, in the vicinity of the Chatham Rise, east of New Zealand (Fig. 1a). We used a Lagrangian experimental array to conduct *in situ* incubations (e.g., ^14^C primary production rate measurements, grazing dilution experiments), which consisted of surface floats for flotation and an iridium-enabled float for satellite tracking of the array. A plastic-coated wire connected the float to a 3×1-m holey-sock drogue centered at 15-m depth to ensure that the Lagrangian array tracked the mixed layer. Mesh bags were placed to span the depth of the euphotic zone, adjusted for each experimental cycle based on the CTD fluorescence profile, and contained experimental rate measurement bottles at six depths within the euphotic zone deployed for 24 hours. A surface-tethered, free-drifting particle interceptor trap (PIT) array was deployed in proximity to the *in situ* array, equipped with a surface Iridium beacon and light flasher, surface and subsurface floats, 10 m-long bungies (to reduce wave action effects) and holey-sock drogue at 15 m to ensure the trap arrays followed the same water parcel as the *in situ* drifting array. A total of five water parcels (experimental cycles) were sampled, for periods ranging from 3-7.5 days (Main text Table 1).

### Physical oceanography

Profiles of temperature, salinity, dissolved oxygen, and photosynthetically active radiation (PAR) were provided by a Seabird (SBE 911plus) Conductivity Temperature – Depth (CTD) with PAR sensor attached to a rosette frame with 24 10L Niskin bottles for water collection. Samples for sensor calibration (salinity and dissolved oxygen) were taken throughout the voyage. Nutrient samples were taken from the Niskin bottles to measure the dissolved inorganic nitrate, ammonium, phosphate, and silicate concentrations at selected depths through the water column.

### Phytoplankton biomass, community composition, and physiological status

The phytoplankton community was sampled from CTD water bottle samples by assessing abundance (flow cytometry, microscopy, and FlowCAM), chlorophyll *a* (chl *a*, total and size-fractionated at 20 μm, 2 μm and 0.2 μm) using fluorometric methods, and chemotaxonomic pigments (including all chlorophyll types, accessory and photoprotective pigments) using High Performance Liquid Chromatography (HPLC). The eukaryotic community within the euphotic zone was assessed using DNA meta-barcoding of the 18S rDNA, from water samples collected during CTD casts.

Phytoplankton physiology was evaluated using a Mini-Fire fast repetition rate fluorometer (FRRF). Fv/Fm and reoxidation of Qa-, the first quinone acceptor of PSII) were evaluated using the MiniFIRe, which provided indications of phytoplankton stress most likely caused by iron-limitation^37^.

#### ^14^C Net primary production (NPP)

Net primary production (NPP) was assessed using ^14^C assays^38^, with 24-hour incubations that integrated respiration/production balance over the dark and light periods of the diel cycle.

Seawater samples (1.3 L) were collected into an acid-rinsed polycarbonate bottle from pre-dawn CTD casts (~0200 h each day of each cycle) at six depths spanning the euphotic zone. The bottles were then spiked with 0.1 mCi ^14^C-bicarbonate (DHI, Denmark or Perkin-Elmer, USA) before triplicate controls on ethanolamine were taken to quantify initial radioactivity at each depth incubation. After gentle mixing, the ‘hot’ 1.3 L was dispensed into three light and one dark bottles (320mL acid-cleaned polycarbonate) that were incubated *in situ* on the free-drifting array. After recovery, the entire content of the bottles were filtered onto 0.2-μm pore-size 25-mm polycarbonate filters and kept frozen until analysis. Once on land, filters were acidified with 200 μL 0.5 N HCl, Hi Safe 3 liquid scintillation cocktail was added and disintegrations per minute were then determined using a scintillation counter following procedures described elsewhere^39^.

#### Phytoplankton growth and microzooplankton grazing

Rates of phytoplankton growth and microzooplankton grazing were assessed daily with the dilution technique^40^, following the two treatment approach^41^, at six depths within the euphotic zone. We implemented this mini-dilution approach to generate vertically resolved growth and grazing rates, but also conducted a full dilution experiment on the last day of each of the cycles (n = 5) to test linearity assumptions of the method. Seawater collected with the Niskin bottles attached to the CTD rosette at 0200 h was used to fill a pair of 2.2-L polycarbonate bottles (100%, B and C) while a third bottle (A) was filled with 25% whole seawater diluted with 0.2-μm filtered seawater obtained immediately before by gravity filtration using an Acropak filter cartridge (Pall) directly from the same Niskin bottle. Nutrients (final concentrations in 2.2L bottles; nitrate 0.18 μM, ammonium 4.16 μM, phosphate 15.08, silicate 44.2 μM, and vitamins) were added to bottles A and B in order to ensure the assumption that the same phytoplankton intrinsic growth rate was occurring in WSW and FSW bottles despite dilution^39^. Bottles were then incubated *in situ* at the same six depths of collection using a drifting array. Rates were calculated from changes in Chl *a* concentration and picophytoplankton abundance between the beginning and end of the experiment assuming exponential growth of phytoplankton.

Microzooplankton grazing rate was estimated from: m μ (k_A_ – k_B_)/(1-x) where k_A_ and k_B_ are the observed net rates of change of chl *a* in bottles A and B, respectively, and x is the fraction of whole seawater in the diluted bottle A (0.25). Phytoplankton growth rate was estimated from: ε = m + k_C_ Photoacclimation effects were corrected from changes in cell chl *a* fluorescence estimated by flow cytometry during incubations as a proxy of cell chl *a* content^42^. These include estimating the photoacclimation index (Phi) from changes in FL3: FSC and calculating an average value from Phi index obtained for pico- and nanoeukaryotic populations weighted by their biomass contribution.

### Pigments (water column, dilution and zooplankton)

Chl *a* and phaeopigments (for water column biota, dilution experiments and zooplankton gut contents) were determined using a Turner 10AU fluorometer with chl *a* filter sets, using the acidification *in vitro* approach^43^. Seawater samples were filtered onto 25mm GF/F filters and flash frozen. Subsequently filters were extracted in 7mL of 90% acetone for 24 h at −20°C. After extraction, samples were brought to room temperature (in the dark), and readings taken with the fluorometer before and after acidification (samples acidified with the addition of 50μL of 10% HCl), to determine chl *a* and phaeopigments. For zooplankton and salp gut pigment determinations, the organisms or their guts were placed in a 15mL Falcon tube containing 6mL of 90% ice-cold acetone, sonicated for 18s, and allowed to extract for 4-20 h. After this period, samples were centrifuged at 3000g for 5 minutes, and chl *a* and phaeopigments were measured in the fluorometer. No correction for pigment destruction was applied^44^.

Chemotaxonomic pigments of phytoplankton in the water column were filtered onto GF/F filters (2L), then immediately flash-frozen in liquid nitrogen, and shipped frozen to Instituto Español de Oceanografía, Centro Oceanográfico de Gijón, where they were extracted for HPLC analyses following established protocols^45^.

### 16S DNA metabarcoding

Bacterial community composition samples were collected from a minimum of six depths, a minimum of three times per cycle. Sample collection consisted of filtering 0.5-1.5L of seawater through a 0.2μm 47mm Millipore filter. The filter was flash-frozen in liquid nitrogen and stored at −80°C until processing. DNA was extracted separately from each filter using a PowerSoil DNA Isolation Kit (Mo Bio, Carlsbad, CA, United States). The manufacturer’s protocol was modified to use a Geno/Grinder for 2 × 15 s instead of vortexing for 10 minutes. DNA concentration was measured using a Nanodrop Spectrophotometer and then a Qubit™ DNA HS Assay Kit, both from Thermo Fisher Scientific. 16S rRNA gene amplicon sequencing was carried out using the Earth Microbiome Project barcoded primer set and conditions^46^. All amplicons (independent replicates) were run on an Illumina HiSeq 151bp x2 run. Amplicon sequence variants (ASVs) were then resolved at single nucleotide resolution using the dada2 pipeline^47^. SILVA release 132 database^48^ was used to assign taxonomy to the sequences. The phyloseq^49^ package in R (R Core Team, 2019) was used for sequence read counts, taxonomic assignments and associated metadata. Sequence reads were randomly rarified to an even depth of 14900 reads per sample prior to analyses. Bray Curtis-based PERMANOVAs using the Adonis function in the vegan package^50^ were used to determine whether the communities were significantly different between water masses and salp blooms.

### 18S DNA metabarcoding of water column and PIT samples

Samples from the water column were collected from six depths within the euphotic zone, filtering 1.4-2.4L per depth. Samples from PITs were collected from four depths (Fig. 2a), and typically contents of one full formalin-preserved sediment trap tube were filtered per sample. Sample collection consisted of filtering contents through 47 mm diameter polycarbonate filter for water column samples (0.2 μm pore size) and serially size-fractionated for PITs samples with 20μm and 0.2 μm pore size filters (Poretics). The filter was flash-frozen in liquid nitrogen and stored at −80°C until processing. DNA was extracted using DNeasy mini kit (Qiagen, Germany) - ‘Qiagen DNA easy Blood and tissue. PCR conditions followed a modified protocol ^51^. Each 50μL reaction included 25 pmol of each primer (V4F_Illumina – 5’ CCAGCASCYGCGGTAATTCC 3’, V4AZig_Illumina – 5’ACTTTCGTTCTTGATYRATGA 3’), 1X KAPA HiFi HotStart ReadyMix (KAPA Biosystems), and 10-50ng of template DNA, with a thermal profile of 95°C for 3 minutes, followed by 10 cycles of 98°C for 10 s, 44°C for 20 s and 72°C for 15 s, followed by 15 cycles of 98°C for 10 s, 62°C for 20 s and 72°C for 15 s, with a final extension of 72°C for 7 minutes. Products were visualized on an agarose gel and successful amplifications were submitted for further adapter ligation and indexing, prior to sequencing on an Illumina MiSeq. Bioinformatics were conducted using dada2^47^ and phyloseq^49^ packages, using the PR2 database version 4.12 (https://pr2-database.org/)^52^ for taxonomic assignation. Bray Curtis-based PERMANOVAs using the Adonis function in the vegan package ^50^ were used to determine whether the communities were significantly different.

### Salp and zooplankton abundance and biomass estimation

Double oblique zooplankton net tows from 200 m water depth to the sea-surface were carried out using a 0.7 m-diameter Bongo frame with paired 200-μm mesh nets, equipped with two General Oceanics flow meters to measure the filtered volume and a temperature-depth recorder. Tows were conducted at least twice daily (day and night), with one additional day per cycle of sampling every 2-3 hours for further studies of diel patterns. Salp specimens were identified to species, using the keys in^53–55^, classified into oozooid or blastozooid stages, measured for total length and corrected to oral to atrial length (OAL)^21^. A random subsample (10 specimens, when available) of each species/stage from each tow was taken for determination of chl *a* in salp guts for grazing estimates. For further biomass and chl *a* analyses, *Salpa thompsoni* lengths were divided into 5mm bins (5-140mm), abundance was calculated for each size bin, and biomass was calculated using length-frequency distributions^23^. The zooplankton (non-salp) sample was split twice to obtain size-fractionated estimates of biomass (dry weight) and gut fluorescence (chl *a* for grazing estimates).

### Salp grazing

Salp specimens (typically 10 of each species/stage if abundance allowed) from each tow had their guts excised, and chl *a* and phaeopigment gut content concentrations were measured. A power function was used to fit the size-specific Gpig (chl *a* + phaeo) contents for each tow, allowing the estimation of Gpig for each size bin per tow, and this was multiplied by the abundance in each size bin. Gut passage time (GPT) was calculated using a modified equation, based on^56^ where: GPT(h) = 2.607*ln(OAL, mm) - 2.6. Grazing was estimated as: G (h^−1^) = Gpig /GPT. Daily salp grazing rates were obtained by assuming 14 h of day and 10 h of night, coincident with the times and latitudes at which we sampled these communities during SalpPOOP.

### Passive export flux – Particle Interceptor sediment Traps (PIT)

Surface-tethered, free-drifting cylindrical PITs with an inside diameter of 7 cm and an 8:1 aspect ratio (height: diameter) were deployed at the start of each experimental cycle^57,58^. Traps were topped by a baffle constructed from smaller 1.3 cm-diameter tubes with a similar 8:1 aspect ratio. Cross-frames, holding 12 baffled PITs, were typically deployed at ~30m below the base of the mixed-layer (as estimated by CTD profiles (T, S, density and fluorescence) and at 100m, 300m and 500m below the sea-surface. PITs were filled with SupraPak 0.2-μm cartridge-filtered seawater and then backfilled to a height of one cylinder diameter (approx. 7-8cm) with hypersaline brine solution (50 g L^−1^) either with or without buffered formaldehyde added (0.4% formaldehyde final concentration) depending on intended analysis. Upon recovery, the overlying seawater was gently siphoned off before the samples were filtered through 200-μm mesh. The mesh filters were then examined under a dissecting microscope (20x magnification) and zooplankton “swimmers” were removed manually from each sample and retained in 5% formalin, prior to photographing the >200μm fraction. Both size fractions were then recombined, and samples were either filtered on pre-combusted GF/F for particulate organic carbon and nitrogen (POC, PON), uncombusted GF/F for chl-*a*, and QMA or membrane filters for C:^234^Thp ratios or DNA meta-barcoding (see below). Additional samples were collected for microscopy. POC and PON were determined on acidified (fumed, HCl) samples run on an isotope ratio mass spectrometer at the UC Davis Stable Isotope Facility (USA).

### Salp fecal pellet contribution to PIT fluxes

The >200μm mesh filters for each PIT tube (used for removing zooplankton “swimmers”) were imaged using a Canon 5D Mark II camera with attached 100mm F/2.8 macro lens mounted in a downward-facing macrophotography rig. Images were manually analyzed using Image J to determine morphometric measurements for each large salp fecal pellet (FP). Morphometric measurements were then used to estimate FP volume and carbon content (see below). These pellet flux measurements should be considered a conservative estimate of total salp FP flux as they only include intact pellets that could be identified from the images taken of the filters. Figures were constructed using ggplot2^59^.

### ^238^Uranium-^234^Thorium Disequilibrium export flux estimates

Twice per cycle ^238^U:^234^Th disequilibrium measurements were taken using the standard small volume method^60,61^. Briefly, 4L water samples were collected by CTD rosette. Sample pH was adjusted to <2 with concentrated HNO_3_, spiked with a yield tracer—10 dpm ^230^Th—and shaken vigorously. After 6-12 hours, pH was adjusted to 8-9 with NH3OH. Co-precipitation with KMnO4 and MnCl2 (100μL each at 7.5g L^−1^ and 33g L^−1^, respectively) was conducted 8-12 hours prior to filtration on a QMA filter which was then dried at 45°C and mounted in a Riso sample cup. Beta activity was then measured with a Riso Ultra-Low Background Beta Multi-counter. After a final beta activity measurement >6 half-lives after collection, filters were digested in an 8M HNO3 / 10% H_2_O_2_ solution, spiked with 5 dpm ^229^Th, and sonicated for 20 minutes. Thorium was then selectively isolated by column chromatography (AG-X8), and isotope ratios (^229^Th:^230^Th) were determined by inductively coupled plasma mass spectroscopy (Thermo Element-2 at the National High Magnetic Field Laboratory, Florida, USA) and used to determine ^234^Th yield. Total ^234^Th activities were used in a 1D water column export model^16^ with corrections for turbulent mixing 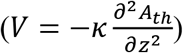 and assuming steady state 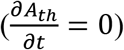. Non-steady state export fluxes were determined for cycle 2 from the rate of change in ^234^Th inventory over the duration of a cycle ^16^. Export fluxes were calculated at the depths corresponding to the PIT deployments, and from which the C:^234^Th ratios were determined (see above). Measurement uncertainties were propagated through all equations.

### Fecal pellet carbon:volume and sinking rates

Salp FPs were collected from live incubations to: *i*) measure carbon:volume FP relationships, and *ii*) measure FP sinking rates. FPs had their length and width measured, and were measured for carbon and nitrogen to create a carbon: volume relationship, which was applied to the pellets imaged in the PITs to estimate the salp FP contribution to export. Carbon content of each FP was determined from a log-log relationship between pellet volume and pellet carbon mass (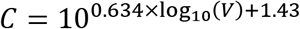, where C is carbon mass in units of μg carbon and V is volume in mm^3^). This relationship was determined from individual fecal pellets that were collected at sea, imaged to determine volume, and analyzed for carbon and nitrogen content on an isotope ratio mass spectrometer.

### Statistical analyses

Bioinformatic analyses were carried out in R. Plots of rates and standing stocks were carried out in MATLAB R2020a, as well as Model I regressions to investigate statistical relationships between export and other explanatory variables.

### Southern Ocean (Polar Frontal Zone) estimates of salp grazing

Salp abundances were downloaded from KRILLBASE, a data rescue and compilation project (https://www.bas.ac.uk/project/krillbase/) that includes the most comprehensive salp abundance dataset for the Southern Ocean (SO), with abundance tows taken from 1926-2016^17^. Net primary production for the same region was estimated using the Vertically-Generalized Production Model (VGPM)^26^ applied to remotely sensed satellite data. Salp grazing rates were estimated for the SO region south of the SAF, based on all relevant data from the SalpPOOP experiment, determined on a carbon-basis. This grazing-vs-abundance relationship assumes that the range of differently sized salps and their relative abundance is representative of the overall general population since previous studies have found grazing-vs-length relationships that are similar to those of our study^62^. Despite the latitudinal/temperature difference between most SO studies and ours off Aotearoa New Zealand, these rates are not substantially different^11,28,56^, suggesting that the results from SalpPOOP for *S. thompsoni* are generally applicable to other regions of the SO.

## Acknowledgments

We thank the captain/crew of the *R/V* Tangaroa and many scientists that assisted on the voyage: Alexia Saint-Macary, Christian Fender, Christopher Ray, David Demory, Florian Lüskow, Lana Young, Matt Walkington, Miguel Méndez Sandín, Morgan Meyers, Natalie Yingling, Rob Stewart, Sadie Mills, Sarah Searson, and Siobhan O’Connor, and the NIWA Vessels Management team for support on land. Thanks to Sven Tobias-Hünefeldt for assistance with DADA2 pipeline analysis. We thank C. Law (NIWA) for comments on the MS.

## Author contributions

M.D. developed the original science questions and cruise plan. M.D., M.R.S., A.G.R., S.D.N., M.P., and K.E.S organized the SalpPOOP voyage and sampling structure. M.D. led the zooplankton/salp sampling and analyses. M.R.S., S.D.N. and T.B.K. led the export flux measurements. A.G.R., K.E.S. and K.S. led the phytoplankton community measurements. A.L.S. and A.G.R. led the eukaryotic metabarcoding measurements. F.D., F.B., and S.E.M. led the bacterial community measurements. M.L. and A.G.R. led the pigment-based phytoplankton community characterization. M.Y.G. and A.G.R. characterized the phytoplankton physiological response. M.P. led the estimation of salp grazing of NPP in the SO. All authors contributed to the development and writing of the manuscript.

## Data and materials availability

Export flux data has been deposited in BCO-DMO (https://www.bco-dmo.org/dataset/813759). All bioinformatics data have been deposited in NCBI (16S data - PRJNA670059 and 18S (CTD and Traps) - PRJNA670061) and code are available on github (https://github.com/Lopesas/TAN1810_Moira_et_al_2020). The remaining datasets presented in the manuscript are available in the tables, or have been deposited in PANGAEA (https://doi.org/10.1594/PANGAEA.928084; https://doi.org/10.1594/PANGAEA.928086 https://doi.org/10.1594/PANGAEA.928087; https://doi.org/10.1594/PANGAEA.928088; https://doi.org/10.1594/PANGAEA.928089; https://doi.org/10.1594/PANGAEA.928092; https://doi.org/10.1594/PANGAEA.928096; https://doi.org/10.1594/PANGAEA.928152; https://doi.org/10.1594/PANGAEA.928170).

