## Supplemental tables and figures for "Salp blooms increase carbon export 5-fold in the Southern Ocean"

### Supplemental Figures

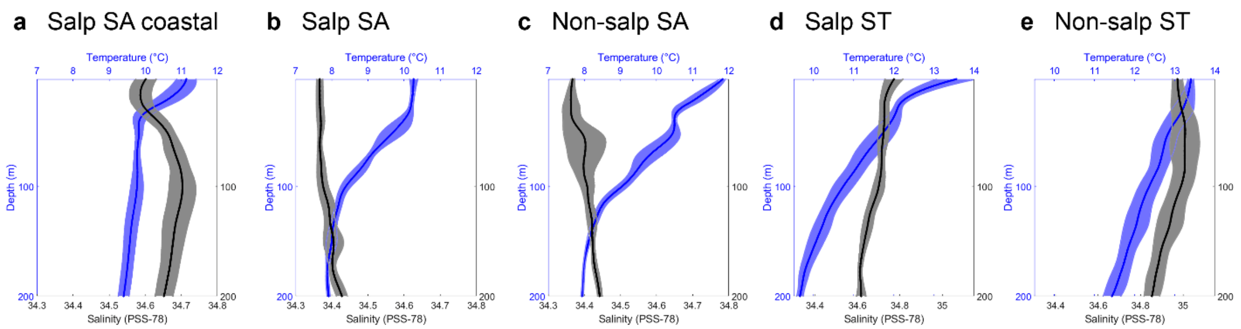


**Supplementary Figure 1. Temperature and salinity profiles (± std) for all five experimental locations**. **a-c**) SA locations, **d-e**) ST areas. Note difference scales in SA compared to ST waters.

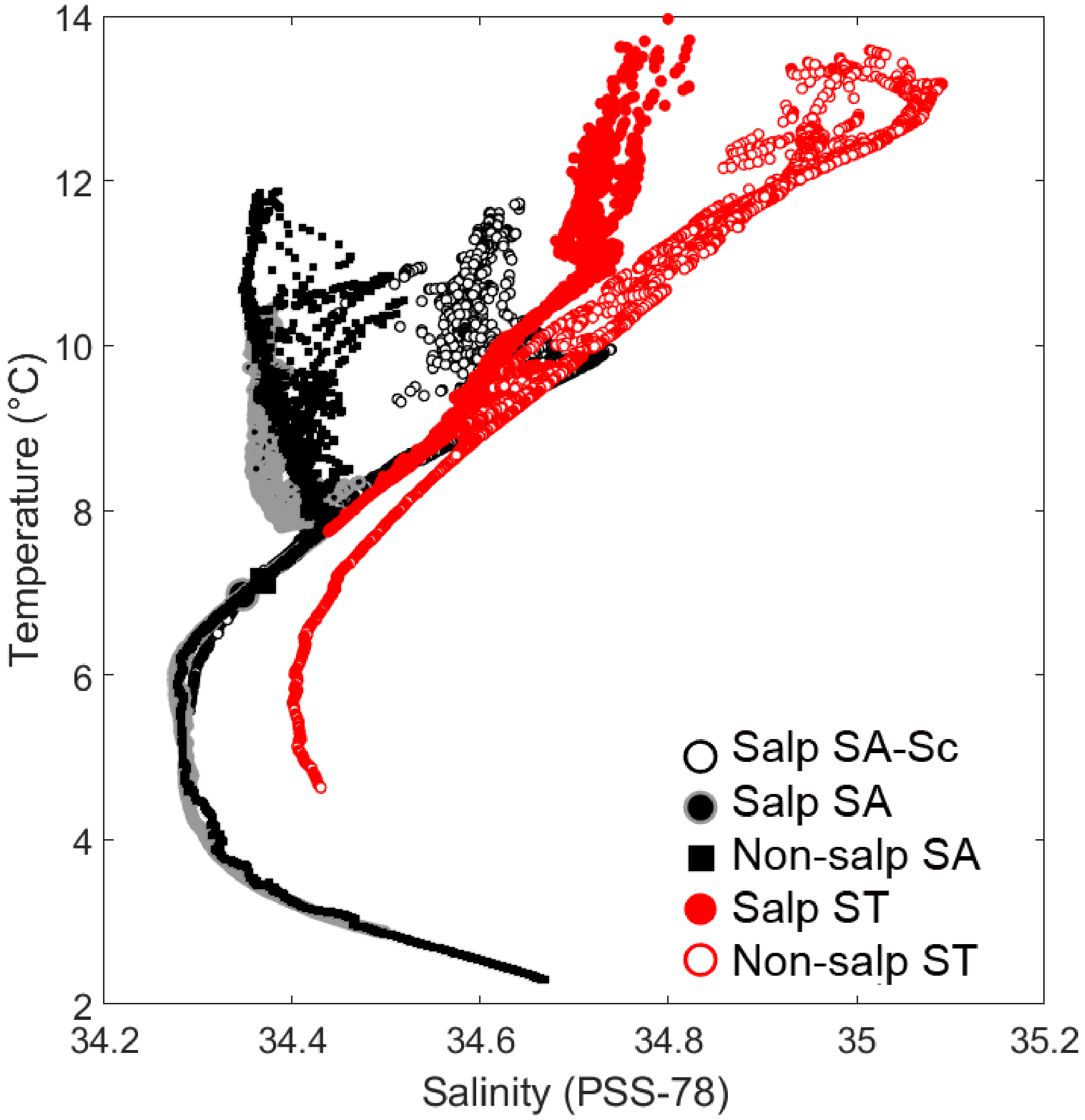


**Supplementary Figure 2. Temperature-Salinity for SalpPOOP cycles**. Black and grey areas represent SA conditions, and red are ST areas.

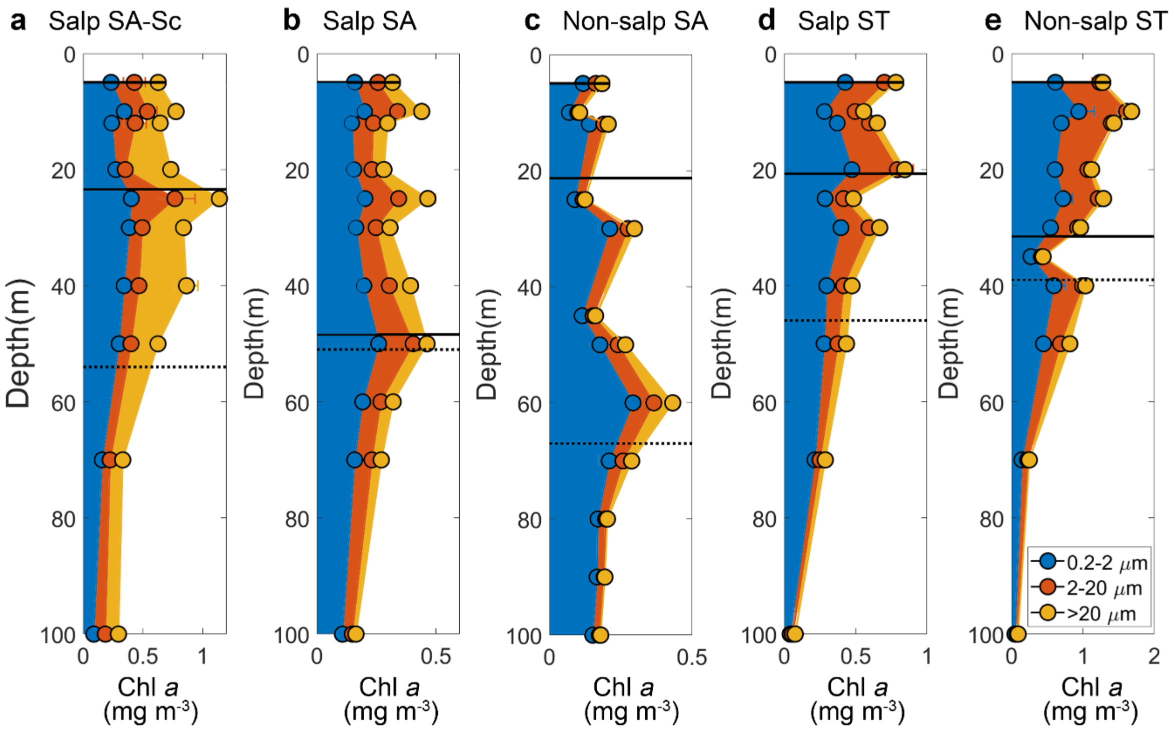


**Supplementary Figure 3. Depth-resolved size-composition (0.2-2µm, 2-20µm, >20µm) of chlorophyll *a*. (a-e)** Corresponds to each experimental cycle. Note different scales on x axis. Mixed layer is denoted by the straight line (shallower depth) and euphotic zone (0.1% PAR) by the dashed line (deeper depth).

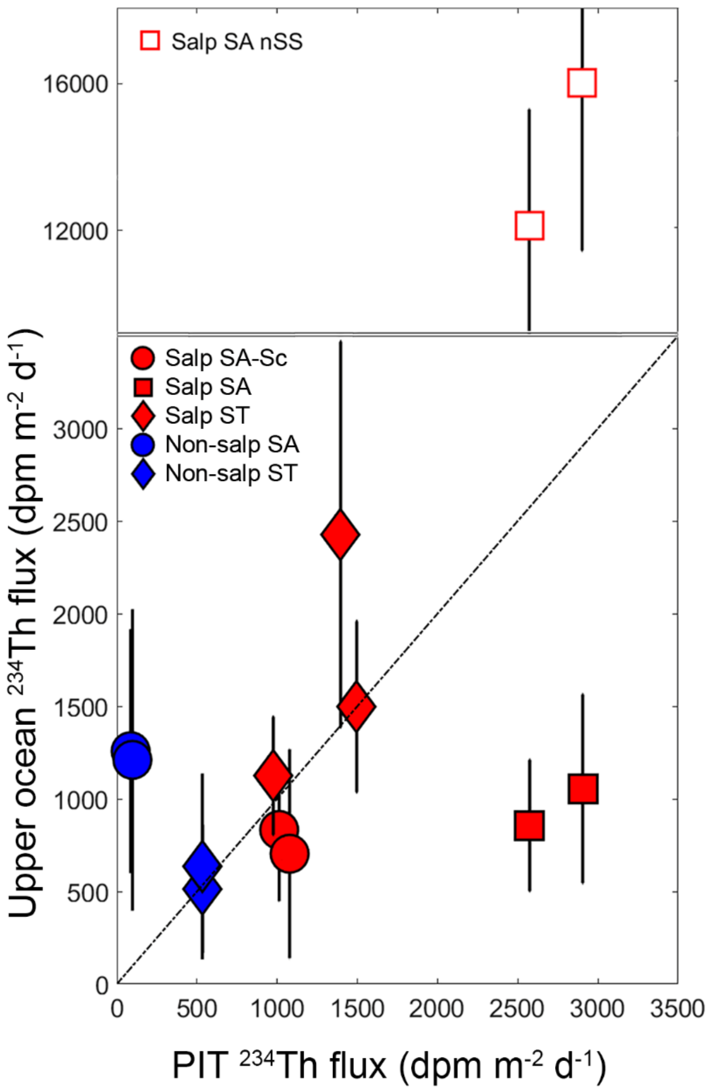


**Supplementary Figure 4. $^{234}$Th flux out of the upper ocean**. X-axis indicates Th-234 flux estimates measured in PITs, while y-axis indicates Th-234 flux estimates using the $^{238}$U:$^{234}$Th disequilibrium method, using water profiles of Th-234 and salinity-derived U-238 estimates, with integration depths matching the depth of the PITs. Filled markers indicate values modelled using steady-state assumptions, open markers (Salp SA only) indicate non steady-state assumptions (see above). Note scale change for non-steady state.

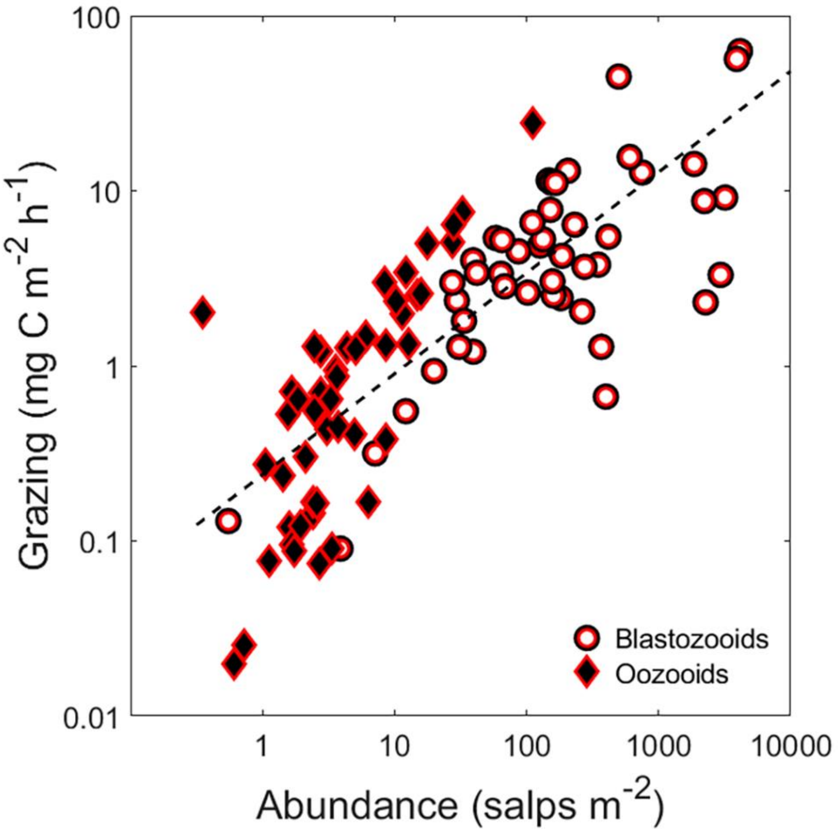


**Supplementary Figure 5. Salp grazing vs abundance.** Linear regression fit (Grazing (mg C $m^{-2}$ $h^{-1}$) = 0.239 * abundance $(m^{-2})^{0.576}$; $r^2 = 0.64$, $p<<0.001$). Data for blastozooids (circles) and oozooids (diamonds) as per the legend; relationship fit is for all data points. Note different log scales on x- and y-axes.

**Supplementary Tables**

| **Pigment** | **Abbreviation** | **Taxonomic significance** |
|---|---|---|
| 19' Butanoyloxyfucoxanthin | 19But | Pelagophytes, Prymnesiophytes |
| 19' Hexanoyloxyfucoxanthin | 19Hex | Prymnesiophytes |
| Alloxanthin | Allox | Cryptophytes |
| Chl *b* | Chl b | Chlorophytes, Prasinophytes |
| Divinyl chl *a* | DVChla | Prochlorophytes |
| Fucoxanthin | Fuco | **Diatoms**, Prymnesiophytes and Pelagophytess |
| Peridinin | Per | Dinoflagellates |
| Prasinoxanthin | Pra | Prasinophytes |
| Zeaxanthin | Zea | Cyanobacteria |

**Supplementary Table 1.** HPLC pigments and main chemotaxonomic phytoplankton group affiliations[1].

| | **Plankton stocks and rates** | **Export 70m** | | **Export 100m** | | **Export 300m** | | **Export 500m** | | **Average** | |
|---|---|---|---|---|---|---|---|---|---|---|---|
| | | **R** | ***p*-value** | **R** | ***p*-value** | **R** | ***p*-** | **R** | ***p*-value** | **R** | ***p*-value** |
| Zooplankton | Zooplankton grazing | 0.14 | 0.79 | 0.37 | 0.47 | 0.14 | 0.80 | 0.58 | 0.31 | 0.18 | 0.73 |
| | Log zooplankton biomass | -0.10 | 0.85 | 0.17 | 0.75 | -0.05 | 0.92 | 0.18 | 0.78 | -0.03 | 0.95 |
| Salps | Salp grazing | 0.26 | 0.62 | 0.51 | 0.30 | 0.22 | 0.67 | 0.76 | 0.13 | 0.31 | 0.54 |
| | Log salp abundance | 0.42 | 0.40 | 0.64 | 0.17 | 0.42 | 0.41 | 0.72 | 0.17 | 0.47 | 0.34 |
| | **Log salp biomass** | 0.66 | 0.15 | **0.78** | **0.07** | 0.65 | 0.16 | 0.77 | 0.13 | 0.69 | 0.13 |
| Chl *a* | Log Surface chl *a* | 0.11 | 0.84 | 0.16 | 0.76 | 0.21 | 0.69 | -0.17 | 0.79 | 0.14 | 0.79 |
| | Log Areal chl *a* | 0.01 | 0.99 | 0.02 | 0.97 | 0.13 | 0.81 | -0.31 | 0.61 | 0.04 | 0.95 |
| Nutrients | Areal nitrate | -0.34 | 0.51 | -0.36 | 0.49 | -0.44 | 0.39 | 0.05 | 0.93 | -0.37 | 0.47 |
| | Areal silicate | -0.47 | 0.34 | -0.40 | 0.43 | -0.54 | 0.27 | 0.05 | 0.93 | -0.49 | 0.33 |
| Phytoplankton | Average Fv/Fm | 0.17 | 0.79 | 0.07 | 0.92 | 0.30 | 0.62 | -0.47 | 0.53 | 0.17 | 0.78 |
| rates | Average PSII reaction center | -0.28 | 0.65 | -0.25 | 0.68 | -0.40 | 0.51 | 0.19 | 0.81 | -0.30 | 0.63 |
| and | Surface NPP | -0.24 | 0.64 | -0.22 | 0.68 | -0.16 | 0.77 | -0.46 | 0.44 | -0.23 | 0.67 |
| physiology | Areal NPP | -0.28 | 0.60 | -0.23 | 0.66 | -0.18 | 0.73 | -0.44 | 0.46 | -0.26 | 0.62 |
| | Biomass accumulation | -0.11 | 0.86 | -0.08 | 0.89 | -0.02 | 0.98 | -0.23 | 0.77 | -0.10 | 0.87 |
| | Chl *a* accumulation | -0.38 | 0.45 | -0.36 | 0.48 | -0.30 | 0.57 | -0.46 | 0.43 | -0.38 | 0.46 |
| | Phytoplankton growth | -0.17 | 0.75 | -0.05 | 0.93 | -0.10 | 0.85 | -0.09 | 0.88 | -0.14 | 0.79 |
| | Microzooplankton grazing | 0.06 | 0.90 | 0.29 | 0.57 | 0.11 | 0.84 | 0.27 | 0.65 | 0.12 | 0.83 |
| 18S DNA | Mamiellophyceae | -0.02 | 0.98 | -0.04 | 0.95 | 0.05 | 0.94 | -0.10 | 0.90 | -0.02 | 0.97 |
| dominant | Dinophyceae | 0.22 | 0.72 | -0.03 | 0.96 | 0.28 | 0.65 | -0.61 | 0.39 | 0.19 | 0.76 |
| phytoplankton | Syndiniales | -0.29 | 0.64 | -0.46 | 0.43 | -0.23 | 0.71 | -0.69 | 0.31 | -0.31 | 0.61 |
| composition | Prymnesiophyceae | 0.01 | 0.99 | -0.08 | 0.89 | -0.08 | 0.90 | 0.06 | 0.94 | -0.02 | 0.97 |
| | Bacillariophyta | 0.01 | 0.98 | 0.27 | 0.65 | -0.03 | 0.96 | 0.64 | 0.36 | 0.05 | 0.94 |
| HPLC | 19' Butanoyloxyfucoxanthin | -0.14 | 0.82 | -0.08 | 0.90 | -0.26 | 0.67 | 0.30 | 0.70 | -0.15 | 0.82 |
| phytoplankton | 19' Hexanoyloxyfucoxanthin | -0.14 | 0.82 | -0.13 | 0.84 | -0.24 | 0.70 | 0.13 | 0.87 | -0.15 | 0.81 |
| composition | Alloxanthin | -0.31 | 0.61 | -0.39 | 0.51 | -0.20 | 0.75 | -0.70 | 0.30 | -0.31 | 0.61 |
| | Chl *b* | 0.72 | 0.17 | 0.44 | 0.46 | 0.74 | 0.15 | -0.10 | 0.90 | 0.68 | 0.21 |
| | Divinyl chl *a* | -0.29 | 0.64 | -0.25 | 0.68 | -0.41 | 0.50 | 0.20 | 0.80 | -0.30 | 0.62 |
| | Fucoxanthin | 0.13 | 0.84 | 0.25 | 0.69 | 0.14 | 0.82 | 0.33 | 0.67 | 0.14 | 0.82 |
| | Peridinin | -0.19 | 0.76 | 0.14 | 0.82 | -0.25 | 0.69 | 0.47 | 0.53 | -0.13 | 0.83 |
| | Prasinoxanthin | 0.53 | 0.36 | 0.27 | 0.66 | 0.65 | 0.24 | -0.71 | 0.29 | 0.51 | 0.38 |
| | Zeaxanthin | -0.43 | 0.47 | -0.47 | 0.42 | -0.51 | 0.38 | -0.13 | 0.87 | -0.45 | 0.44 |

**Supplementary Table 2.** Regression results of plankton stocks and rates as explanatory variables of POC export fluxes at four depths, and average export flux for all five experimental cycles. Note that export at 500m does not include Salp ST because export was not measured that deep at that location. The only correlation that had $p<0.1$ is shown in bold; no variables were significant at $p<0.05$.
